# No SARS-CoV-2 cross-neutralization by intravenous immunoglobulins produced from plasma collected before the 2020 pandemic

**DOI:** 10.1101/2020.07.30.228213

**Authors:** Julia Schwaiger, Michael Karbiener, Claudia Aberham, Maria R. Farcet, Thomas R. Kreil

## Abstract

The 2020 SARS-CoV-2 pandemic is caused by a zoonotic coronavirus transmitted to humans, similar to earlier events. Whether the other, seasonally circulating coronaviruses induce cross-reactive, potentially even cross-neutralizing antibodies to the new species in humans is unclear. The question is of particular relevance for people with immune deficiencies, as their health depends on treatment with immunoglobulin preparations that need to contain neutralizing antibodies against the pathogens in their environment. Testing 54 IVIG preparations, produced from plasma collected in Europe and the US, highly potent neutralization of a seasonal coronavirus was confirmed, yet no cross-neutralization of the new SARS-CoV-2 was seen.

**SUMMARY:** IVIG products manufactured from pre-pandemic plasma do not neutralize SARS-CoV-2 but contain high neutralizing titers for seasonal coronavirus hCoV-229E.

## Background

Intravenous immunoglobulins (IVIG) are produced from thousands of pooled plasma donations, and thus contain a wide variety of antibodies the donors have generated against infectious disease agents. IVIG can therefore protect people with immune deficiencies (PID) against circulating bacterial and viral infections.

When a new pathogen emerges, antibodies to the new agent only become detectable in IVIG preparations after a certain proportion of plasma donors have contracted the infection, and successfully recovered from it. An even higher number of convalescent plasma donors is needed to result in neutralizing antibody (nAb) levels high enough for the resulting IVIG to afford protection against the new infectious agent. After the arrival of West Nile virus (WNV) in the US, for example, it took several years before the prevalence of WNV nAbs reached approximately 0.5% in the plasma donor community, at which point they became detectable in IVIG lots produced from plasma donated in the US. Thereafter, a significant proportion of IVIG lots even reached *in vivo* protective levels [1].

Severe acute respiratory syndrome coronavirus-2 (SARS-CoV-2) spread around the globe at unprecedented speed, infecting millions of people already within the first 6 months of circulation. The virus belongs to the *Coronaviridae* family of viruses which contains several species of importance for human health. The human coronaviruses (hCoVs) 229E, NL63, OC43, and HKU1 are in circulation as seasonal respiratory viruses that mostly cause self-limiting and mild infections but which can also lead to pneumonia and bronchiolitis [2–4], whereas the Middle East respiratory syndrome coronavirus (MERS-CoV) causes a prolonged outbreak mainly limited to the Arabian Peninsula, and the SARS-CoV and SARS-CoV-2 viruses have established global transmissions chains – the latter three associated with significant human mortality.

Due to the widespread and long-term circulation of hCoVs, and the pooling of plasma from thousands of donors for every lot of IVIG, it was shown that IVIGs contain significant levels of hCoV nAb levels, as was shown for example for hCoV-NL63 [5]. Whether these hCoV antibodies cross-react with or even neutralize the related SARS-CoV-2 has not been fully elucidated. To date, antibody binding assays have shown some cross-reactivity between different hCoVs and SARS-CoV-2, yet functional and therefore clinically more relevant virus neutralization assays have shown no or only very low levels of cross-reactive antibodies [6–8]. The question is of significant clinical relevance, as SARS-CoV-2 cross-neutralizing antibodies in IVIGs, if they were present, might afford some protection to PIDs, and may even represent a treatment option for COVID-19 patients.

The current study tested a representative number of IVIG lots for nAbs against SARS-CoV-2 and the already longer circulating hCoV-229E, to establish clarity about cross-neutralization of the pandemic virus by antibodies induced by earlier circulating seasonal coronaviruses. In addition, results from the ongoing monitoring of the plasma donor community for the development of SARS-COV-2 antibodies are presented.

## Methods

### IVIG preparations

A total of 54 IVIG lots fractionated from plasma collected prior to the circulation of SARS-CoV-2 were analyzed. The IVIG lots were manufactured from plasma either donated by plasmapheresis (source, S), or recovered from whole blood donations (recovered, R), in the US (Gammagard Liquid; Baxter Healthcare Corp., Westlake Village, CA; N = 30), or central Europe (Austria, Germany, Czech Republic (KIOVIG; Baxter AG, Vienna, Austria; N = 24). These 54 IVIG lots were tested in two independent experiments for nAbs (one EU lot in single assay, due to volume constraints).

### Human plasma samples

Two pre-pandemic plasma donations were collected in April and May 2019, respectively by BioLife Austria.

Human plasma pool samples were generated by combination of 6 individual donations obtained in the same calendar week (CW) at Austrian plasma donation centers (BioLife). The pools were assembled from donations collected in CW 13 (n=40), CW 14 (n=80), CW 15 (n=80), CW 16 (n=80), CW 20 (n=100), CW 24 (n=80), and CW 28 (n=100) of 2020; i.e. during the SARS-CoV-2 pandemic. Cumulative incidence of COVID-19 was calculated from data provided by the Austrian Federal Ministry for Social Affairs, Health, Care and Consumer Protection (www.sozialministerium.at) and Statistics Austria (www.statistik.at).

### Detection of SARS-CoV-2 neutralizing antibodies

SARS-CoV-2 nAb titers were determined in IVIG and human plasma samples that were used undiluted, or pre-diluted with cell culture medium 1:4, 1:5, 1:10 or 1:20 depending on sample amount available, and then serially diluted in 2-fold steps. Equal volumes of sample dilutions were mixed with virus stock at 10^3.0^ tissue culture infectious doses 50% per milliliter (TCID50/mL) SARS-CoV-2 (strain “BavPat1/2020”, kindly provided by C. Drosten and V. Corman, Charité Berlin, Germany) and incubated for 150min +/-15min, before titration on Vero cells (Cat. no. 84113001, ECACC, Porton Down, Salisbury, UK) in eight-fold replicates per dilution. The virus-induced cytopathic effect was determined after 5-7 days of incubation. The reciprocal sample dilution resulting in 50% virus neutralization (NT50) was determined using the Spearman-Kärber formula, and the calculated neutralization titer for 50% of the wells reported as 1:X. The detection limits were as follows: <1:0.8 for undiluted, <1:3.1 for 1:4 prediluted, <1:3.9 for 1:5 prediluted, <1:7.7 for 1:10 prediluted and <1:15.4 for 1:20 prediluted IVIG.

The neutralization assay (μNT) included several validity criteria, i.e. confirmatory titration of input virus infectivity, cell viability, and neutralization testing of an internal reference standard, all of which had to comply with defined ranges.

### Detection of hCoV-229E neutralizing antibodies

The neutralization assay for hCoV-229E antibodies is essentially identical to the SARS-CoV-2, where samples were used undiluted or pre-diluted, then serially diluted in 2-fold steps and mixed 1:2 with 10^3.0^ TCID50/mL hCoV-229E (ATCC, Cat. no. VR-740, Rockville, MD), incubated and titrated on MRC-5 cells (ATCC, Cat. no. CCL-171, Rockville, MD). The virus-induced cytopathic effect was determined after 7-9 days of incubation.

### Graphs and statistical analysis

Graphical illustration and statistical analysis (paired t-tests) were done using GraphPad Prism v8.1.1 software (San Diego, CA).

## Results

The validity and reliability of the SARS-CoV-2 neutralization assay used in the current study was demonstrated by the analysis of 100 convalescent plasma donations from PCR-confirmed SARS-CoV-2 cases [9]. As controls, two plasma samples collected before the emergence of SARS-CoV-2 did not contain SARS-CoV-2 nAbs, as expected, yet neutralized hCoV-229E with NT50 values of 1:43 and 1:26, respectively.

### Testing of IVIG

SARS-CoV-2 nAb titers were below the limit of detection for all 54 IVIG lots tested, irrespective of geographic origin of the plasma (EU vs. US) and plasma collection modality (recovered vs. source) (Figure 1A).

**Figure 1.**
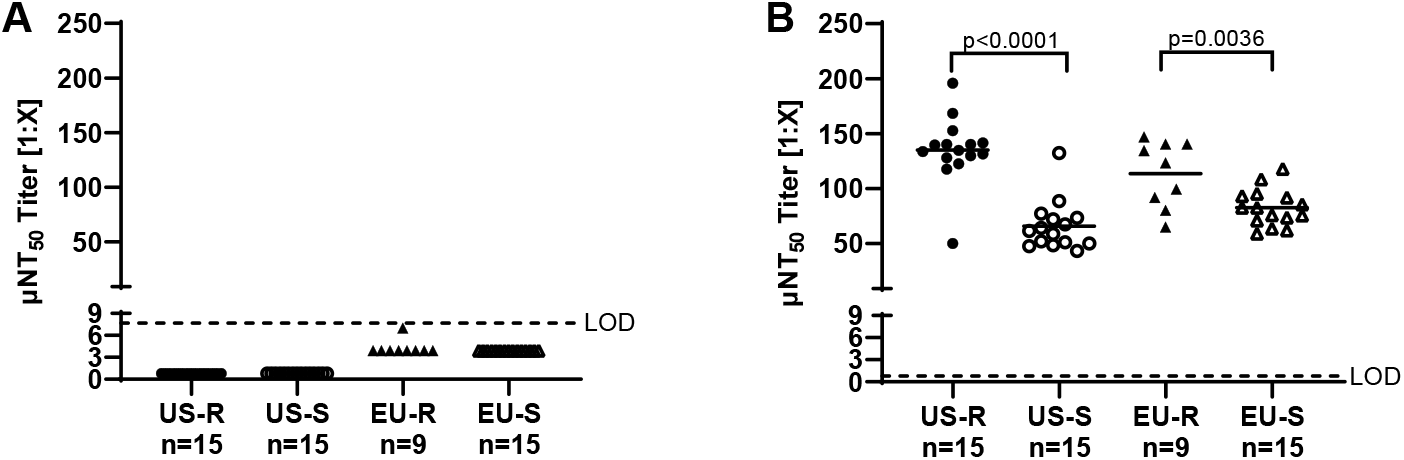
Coronavirus neutralizing antibody titers in intravenous immunoglobulin (IVIG) lots (n=54) against **A.** SARS-CoV-2 and **B.** hCoV-229E. The IVIG lots were manufactured from plasma either donated by plasmapheresis (source, S), or recovered from whole blood donations (recovered, R), in the US or central Europe. Each dot represents the mean of 2 independent experiments (except in panel B: from 1 EU-R IVIG lot the titer of a single determination is shown). The lines represent the median in each group. Paired t-tests were used for determination of significance. LOD: limit of detection;

In contrast, hCoV-229E nAb titers between 1:43 and 1:196 (mean=1:98) were measured for the 54 IVIG lots tested (Figure 1B). IVIG lots produced from recovered plasma contained significantly higher levels of nAb to hCoV-229E as compared to IVIG lots produced from source plasma, independent of the geography of origin. A significant, although quantitatively minor, difference was also found between IVIG lots manufactured from source plasma collected in either the US or the EU.

### Testing of human plasma

To evaluate the potential development of SARS-CoV-2 antibodies in the plasma donor community, samples of plasma pools of 6 donations each were tested. The use of pools enabled to test a high amount of plasma donations for SARS-CoV-2 nAbs with the BSL-3 functional assay. As the mean SARS-CoV-2 μNT50 of plasma donations is rather high (approx. 1:230 [9]) even the nAbs of only one positive sample within a pool are detectable in this assay.

Testing a total of 560 plasma pools of 6 donations each, in total reflective of 3,360 plasma donations, from CW 13 until CW 28, 2020 revealed that most of these pools had SARS-CoV-2 μNT titers below the limit of detection (Table 1). The first pool with detectable nAbs to SARS-CoV-2 was detected in CW 14. Further positive pools were found in CW 15, CW 16, CW24 and CW 28. Up to 7% of the tested pools showed nAbs to SARS-CoV-2, which indicates that up to 1.17% of the plasma donors were positive for SARS-CoV-2 nAbs at a cumulative incidence of COVID-19 in Austria of 0.21% (Table 1).

**Table 1.**
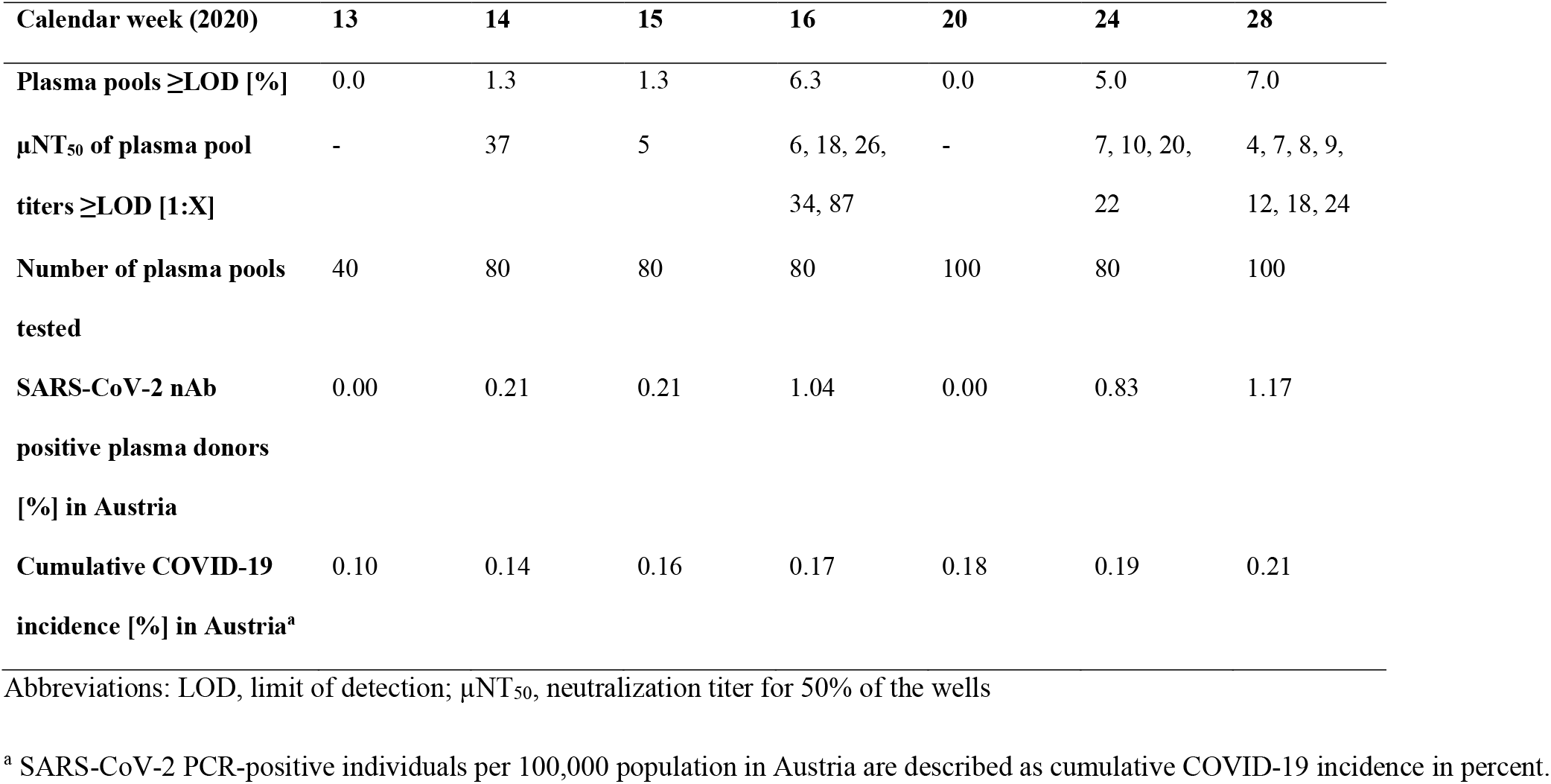
SARS-CoV-2 neutralizing antibodies in tested plasma pools.

## Discussion

Our analysis of IVIG lots revealed high nAb titers to hCoV-229E (Figure 1B). It is interesting to note that in IVIG lots produced from recovered plasma significantly higher titers to hCoV-229E were found compared to IVIG lots produced from source plasma, independent of plasma origin. Recovered plasma is usually obtained from elder donors. A trend towards increasing probability of hCoV-229E infections with age was detected in the Scottish [3] and US [4] population. Furthermore, higher hCoV-229E specific nAb titers were found in older study participants (60-85 years) compared to younger ones (21-40 years) [10]. Thus, a higher infection rate with hCoV-229E in elder people and consequently the generation of neutralizing hCoV-229E-specific antibodies in the elder plasma donors would explain the higher hCoV-229E titers found in IVIG lots produced from recovered plasma.

Low levels of cross-neutralizing Abs have earlier been demonstrated between specific pairs of coronaviruses, specifically between SARS-CoV-2 sera and SARS-CoV [6] and SARS-CoV sera and MERS [11].

In clear contrast, testing the same lots of IVIG that were shown to contain high hCoV-229E nAb titers with a highly specific SARS-CoV-2 μNT did not detect SARS-CoV-2 specific nAbs. The finding confirms that the existing hCoV-229E-specfic nAbs, as well as the presumably present nAbs against the other seasonal hCoVs, have no cross-neutralizing capacity to SARS-CoV-2. These results are entirely consistent with the absence of SARS-CoV-2 antibodies in the plasma used for production of these IVIG lots, as this plasma was donated well before the start of the SARS-CoV-2 pandemic.

Surprisingly, Diez et. al. published the detection of cross-reactive [12] as well as cross-neutralizing Abs in 5 of 6 tested IVIG lots (Flebogamma and Gamunex C) [13]. This is in contrast to an earlier study which tested 21 IVIG lots (9 Gamunex C, 10 Gammagard Liquid, 2 other) with a SARS-CoV-2-specific ELISA (RBD or spike protein) that was shown to correlate well with a neutralization test, and revealed the absence of cross-reactive antibodies [14], as well as the current study. It is worth noting that the two different methods used by Diez et.al. showed up to 5.000-fold different neutralization titers for the same IVIG preparation (G1, compare Fig. 2 and 3 of [13]).

As two experimentally robust studies have not found SARS-CoV-2 nAbs in IVIG lots produced from pre-pandemic plasma, currently available IVIGs cannot be expected to afford protection from SARS-CoV-2 infection.

With increasing numbers of human infections, also in the plasma donor community, it is interesting to follow the development of antibodies against SARS-CoV-2 in plasma donations and, after the several months production cycle time between plasma donation and IVIG lot release, also in IVIG lots, a longitudinal study currently in progress. In this context it is noteworthy that the mean nAb titers induced by WNV [1] and SARS-CoV-2 [9] infection are of similar magnitude. After the emergence of WNV in the US, nAb titers became detectable in IVIGs after approx. 0.5% of the population had gone through and recovered from the infection. In Austria, testing of plasma pool samples indicated that up to 1.17% of plasma donors were positive for SARS-CoV-2 nAbs. Based on the current number of reported SARS-CoV-2 infections in the US (~4.3 million per July 30, 2020; www.cdc.gov), and an estimated rate of >40% asymptomatic infections [15], more than 7.2 million people in the US could have been infected already, i.e. 2.2% of the approx. 330 million population. Based on these facts, the detection of SARS-CoV-2 nAbs in IVIG lots produced from US plasma, the major source for fractionation, is expected within the next few months.

Another, more immediately available possibility for the treatment of COVID-19 is production of a hyper-IVIG from the plasma of COVID-19 convalescent donors (CoVIg-19), a developmental work currently under way through a large alliance of plasma stakeholders (https://www.covig-19plasmaalliance.org).

## Acknowledgments

The contributions of the entire Global Pathogen Safety team, most notably Simone Knotzer, Melanie Graf, Jasmin de Silva, Julius Segui (neutralization assays), Veronika Sulzer, Sabrina Brandtner (cell culture) as well as Eva Ha and Alexandra Schlapschy-Danzinger (virus culture) are gratefully acknowledged. The team of Plasma Analytics, Takeda, Vienna prepared the human plasma pool samples. SARS-CoV-2 was sourced via EVAg (supported by the European Community) and kindly provided by Christian Drosten and Victor Corman (Charité Universitätsmedizin, Institute of Virology, Berlin, Germany).

## References

1. Planitzer CB, Modrof J, Yu MY, Kreil TR. West Nile virus infection in plasma of blood and plasma donors, United States. Emerg Infect Dis 2009; 15:1668–70.

2. Greenberg SB. Update on Human Rhinovirus and Coronavirus Infections. Semin Respir Crit Care Med 2016; 37:555–71.

3. Nickbakhsh S, Ho A, Marques DFP, McMenamin J, Gunson RN, Murcia PR. Epidemiology of Seasonal Coronaviruses: Establishing the Context for the Emergence of Coronavirus Disease 2019. J Infect Dis 2020; 222:17–25.

4. Killerby ME, Biggs HM, Haynes A, Dahl RM, Mustaquim D, Gerber SI, Watson JT. Human coronavirus circulation in the United States 2014-2017. J Clin Virol 2018; 101:52–6.

5. Pyrc K, Bosch BJ, Berkhout B, Jebbink MF, Dijkman R, Rottier P, van der Hoek L. Inhibition of human coronavirus NL63 infection at early stages of the replication cycle. Antimicrob Agents Chemother 2006; 50:2000–8.

6. Anderson DE, Tan CW, Chia WN, et al. Lack of cross-neutralization by SARS patient sera towards SARS-CoV-2. Emerg Microbes Infect 2020; 9:900–2.

7. Huang AT, Garcia-Carreras B, Hitchings MDT, et al. A systematic review of antibody mediated immunity to coronaviruses: antibody kinetics, correlates of protection, and association of antibody responses with severity of disease. medRxiv 2020.04.14.20065771 [Preprint]. April 17, 2020 [cited 2020 Jul 30]. Available from: https://doi.org/10.1101/2020.04.14.20065771.

8. Okba NMA, Muller MA, Li W, et al. Severe Acute Respiratory Syndrome Coronavirus 2-Specific Antibody Responses in Coronavirus Disease Patients. Emerg Infect Dis 2020; 26:1478–88.

9. Jungbauer C, Weseslindtner L, Weidner L, Gänsdorfer S, Farcet MR, Gschaider-Reichhart E, Kreil TR. Characterization of 100 sequential SARS-CoV-2 convalescent plasma donations. bioRxiv 2020.06.21.163444 [Preprint]. June 21, 2020 [cited 2020 Jul 30]. Available from: https://doi.org/10.1101/2020.06.21.163444.

10. Gorse GJ, Donovan MM, Patel GB. Antibodies to coronaviruses are higher in older compared with younger adults and binding antibodies are more sensitive than neutralizing antibodies in identifying coronavirus-associated illnesses. J Med Virol 2020; 92:512–7.

11. Chan KH, Chan JF, Tse H, et al. Cross-reactive antibodies in convalescent SARS patients’ sera against the emerging novel human coronavirus EMC (2012) by both immunofluorescent and neutralizing antibody tests. J Infect 2013; 67:130–40.

12. Diez JM, Romero C, Gajardo R. Currently available intravenous immunoglobulin contains antibodies reacting against severe acute respiratory syndrome coronavirus 2 antigens. Immunotherapy 2020; 12:571–6.

13. Diez JM, Romero C, Vergara-Alert J, et. al. Cross-neutralization activity against SARS-CoV-2 is present in currently available intravenous immunoglobulins. bioRxiv 2020.06.19.160879 [Preprint]. June 19, 2020 [cited 2020 Jul 30]. Available from: https://doi.org/10.1101/2020.06.19.160879.

14. Amanat F, Stadlbauer D, Strohmeier S, et al. A serological assay to detect SARS-CoV-2 seroconversion in humans. Nat Med 2020; 26(7):1033–1036

15. Oran DP, Topol EJ. Prevalence of Asymptomatic SARS-CoV-2 Infection: A Narrative Review. Ann Intern Med 2020; M20–3012.

